# Cega: A Single Particle Segmentation Algorithm to Identify Moving Particles in a Noisy System

**DOI:** 10.1101/2020.12.24.424334

**Authors:** Erin M. Masucci, Peter K. Relich, E. Michael Ostap, Erika L. F. Holzbaur, Melike Lakadamyali

## Abstract

Improvements to particle tracking algorithms are required to effectively analyze the motility of biological molecules in complex or noisy systems. A typical single particle tracking (SPT) algorithm detects particle coordinates for trajectory assembly. However, particle detection filters fail for datasets with low signal-to-noise levels. When tracking molecular motors in complex systems, standard techniques often fail to separate the fluorescent signatures of moving particles from background noise. We developed an approach to analyze the motility of kinesin motor proteins moving along the microtubule cytoskeleton of extracted neurons using the Kullback-Leibler (KL) divergence to identify regions where there are significant differences between models of moving particles and background signal. We tested our software on both simulated and experimental data and found a noticeable improvement in SPT capability and a higher identification rate of motors as compared to current methods. This algorithm, called Cega, for ‘find the object’, produces data amenable to conventional blob detection techniques that can then be used to obtain coordinates for downstream SPT processing. We anticipate that this algorithm will be useful for those interested in tracking moving particles in complex in vitro or in vivo environments.

## INTRODUCTION

Developments in fluorescence imaging methods, such as total internal reflection fluorescence (TIRF) microscopy, have revolutionized live cell microscopy, allowing for monitoring of dynamic events in a variety of biological systems. TIRF uses an evanescent wave generated by reflecting a laser beam at a critical angle to illuminate fluorophores within a very small distance (~100 nm) from the surface of a glass coverslip. The high signal-to-noise achieved by this method improves spatiotemporal resolution over conventional imaging techniques, allowing the monitoring of fluorescent single molecules, molecular complexes, and organelles with nanometer precision on sub-second time scales. TIRF microscopy has been widely used to investigate the movement of molecular motors such as kinesin, dynein and myosin, on their corresponding cytoskeletal tracks in purified systems and cells (Belyy & Yildiz, 2014; Pierce et al., 1997; Vale et al., 1996).

There have been substantial efforts to automate the tracking and analysis of the movement of motors and cargos from movies acquired with TIRF microscopy using single particle tracking (SPT) (Chenouard et al., 2009; Jaqaman et al., 2008; Kalaidzidis, 2007; Meijering et al., 2006; Tinevez et al., 2017). SPT describes the set of techniques that select particles of interest and aggregate temporally separated particle coordinates into trajectories. The resulting trajectories are used to study the dynamics of target particles, including processive movements, diffusion, and pausing. Thus, SPT algorithms must be able to select and follow a target with high spatial and temporal fidelity.

The performance of many SPT algorithms are limited by pre-processing steps that remove background signals and noise that can complicate the generation of particle coordinates that are linked into trajectories (Smal & Meijering, 2015). Current tracking software such as u-track (Jaqaman et al., 2008) and TrackMate (Tinevez et al., 2017) have made great strides to assist in analysis of typical *in vitro* systems that use TIRF microscopy to image movement of purified single motors along separated and immobilized tracks in assays where the signal density can be controlled. However, analysis of low signal-to-noise data is difficult if not impossible when using more complex *in vitro* systems, including engineered cytoskeletal bundles and extracted cytoskeletal systems where bidirectional transport occurs. Images and times series acquired from these more complex *in vitro* systems include substantial background fluorescence and nuisance particles that complicate application of SPT. The available software implementations of SPT include limited solutions for particle segmentation (Jaqaman et al., 2008; Tinevez et al., 2017) that fail when challenged with datasets that include high levels of background fluorescence, and these algorithms do not consider removal of nuisance particles. In addition, these algorithms fail to accurately track particles that change direction over time and cannot be used in systems that display bi-directional movement.

Here, we present a filtering method, Cega, or more appropriately ce:ga (Tohono O’odham for find the object (Mathiot, 1973)), for finding and tracking moving fluorescent objects acquired from experiments performed under conditions that include high noise (Figure 1). This method can be substituted into the initial candidate-finding phase of current SPT software and modified for specific dataset needs for more accurate particle detection. Our method was developed to analyze data acquired from an EMCCD camera with a back projected pixel size that slightly under samples the Nyquist rate for the Point Spread Function of a diffraction limited single molecule. First, we calibrated the camera to properly parameterize the noise statistics. Then, we focused on removing nuisance background signal using Cega. We used multiplicative noise statistics to generate two contrasting models that can then be used to augment the signal of moving motors while suppressing the signal from background and nuisance particles. We tested this model with both simulated and experimental data and found that it performed better than current algorithms in tracking molecular motors moving through complex cytoskeletal arrays that contribute high noise levels. The results suggest that this method provides a substantial improvement for low signal-to-noise tracking applications.

**Figure 1.**
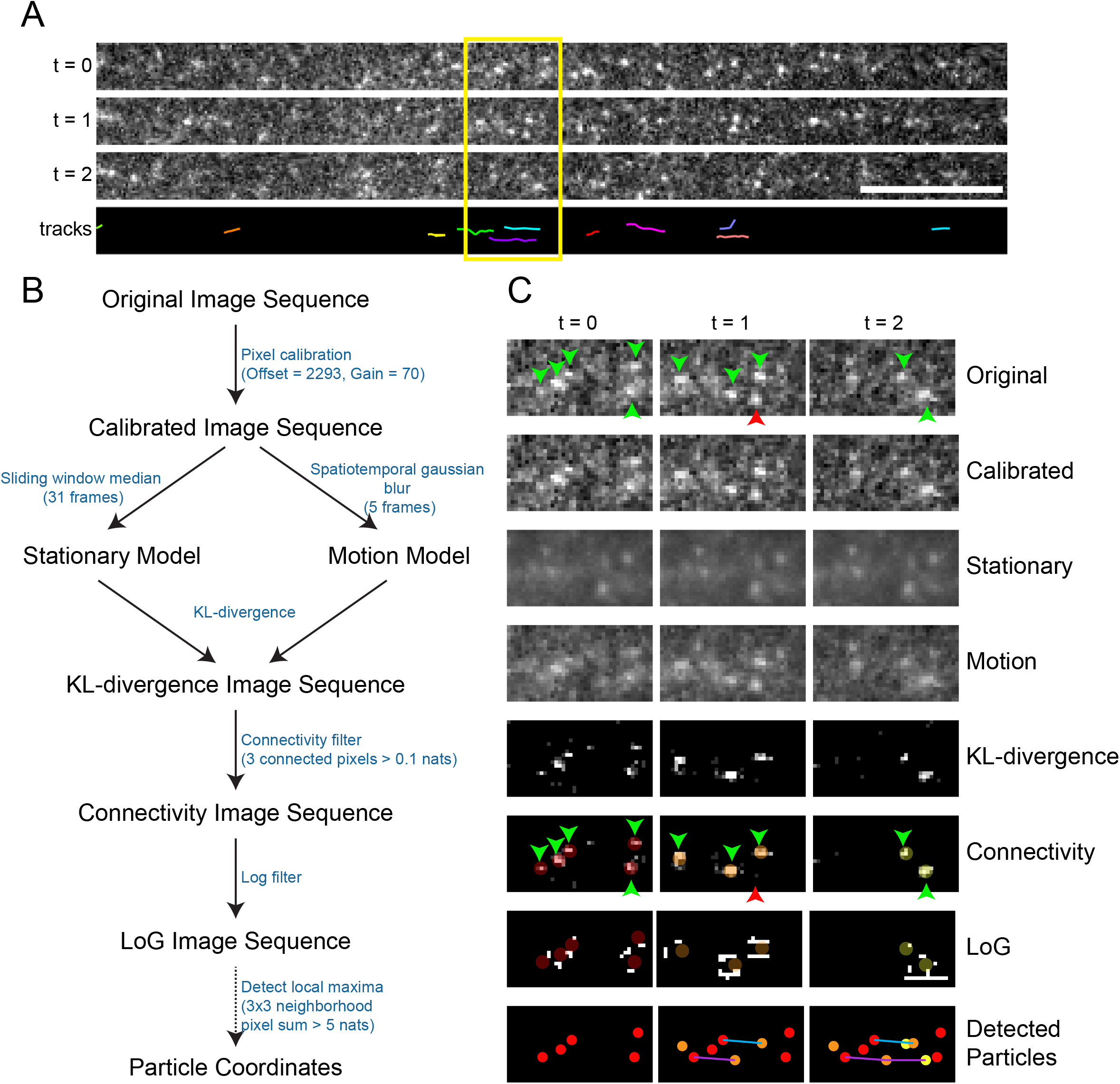
Cega workflow. A) Example time sequence taken from image sequence of kinesin motors moving within an axonal compartment, as well as corresponding tracks calculated after Cega detection. B) Diagram of Cega steps leading to spot detection. Font between each step indicates values used to process axonal and dendritic data. C) Image sequence of timepoints in A after processing with each of Cega’s steps. Image stills represent how data is manipulated at each step. Green arrows indicate moving spots, while red arrows indicate positions of stationary spots. Dim colored spots in the KL-divergence images represent locations of spot detection. Spots in the LoG images represent detected spots, colored by appearance over time. Spots corresponding to the same track are connected with the same colored lines.

## RESULTS

### Computational Strategy

Here we outline our computational approach to detect and track single fluorescent particles that move bidirectionally from data sets with low signal-to-noise. We first provide an overview, then discuss implementation of this strategy to analyze single molecule data sets of kinesin motors moving within complex arrays, and conclude with a comparison of Cega to other computational approaches currently available. The normalization procedures and algorithms we developed are described in more detail in the Methods section, and the corresponding computer code is available online at https://github.com/prelich/Cega.

#### Camera Calibration

Before application of any algorithm, such as Cega, to detect and identify particles of interest in a time sequence, the detector being used to collect the primary data must be calibrated to ensure pixel intensities are appropriately accounted for when analyzed. Individual frames of time sequences obtained from cameras and optical sensors typically have noise statistics that are poorly described by a Poisson distribution, as a result of damaged detectors as well as additive and multiplicative noise that accompanies the conversions of photons to digital signals. These factors complicate the use of algorithms to eliminate noise since the data cannot be assumed to have Poisson statistics, and post-processing data cannot be thresholded reliably (Mortensen et al., 2010; Mortensen & Flyvbjerg, 2016). Thus, camera calibration is required before using Cega to linearly transform the measured pixel intensities of a series of time frames to generate Poisson-like statistics. Specifically, a stationary object with a constant rate of photon emissions acquired over several frames will register pixel intensities that satisfy the following relation:

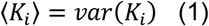

where, given a constant emission of photons at a pixel *i*, the time averaged intensity of the calibrated pixel, < K_i_ >, is equal to the temporal variance of that calibrated pixel, *var*(*K*_*i*_). A Poisson-like distribution is a non-integer analog of the Poisson distribution with units of ‘effective photons’. For our Poisson-like model the probability of observing some positive real number K given an expected value μ is defined as:

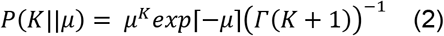

The digital signal recovered from a scientific camera will be reported in Arbitrary Digital Units (ADUs) because pixel statistics are dependent on the camera’s configuration. ADUs are sufficient quantities for relative comparisons or when the follow up analysis treats signal noise as irrelevant or predominantly additive. For algorithms that factor in multiplicative noise, the conversion factors that relate a set of ADUs to effective photons are required quantities. Extensive work on CCD (Young et al., 1998) and sCMOS (Babcock et al., 2019; Huang et al., 2013) sensors has demonstrated that statistical regression with a simplified model can effectively recover Poisson-like statistics for estimators that use multiplicative noise models. Given that a scientific camera produces an output signal *S*_*i*_ at pixel *i* the signal components are defined as:

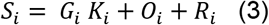

where *K*_*i*_ is the effective photon count, *G*_*i*_ is the multiplicative gain factor on the Poisson-like signal, *O*_*i*_ is the constant mean offset, and R_*i*_ is an unbiased random electronic signal generated from the camera read noise. The purpose of camera calibration is to recover *G*_*i*_, *O*_*i*_, and *var*(*R*_*i*_) and to use these quantities to estimate the effective photons in movies acquired from the calibrated camera. Although the model used to recover *K*_*i*_ is independent of sensor type, calibration techniques will vary among cameras. For EMCCD cameras, pixels are read out in serial so it’s typically assumed that *R*_*i*_ = *R*, *O*_*i*_ = *O, and G*_*i*_ = *G* (see Methods for estimation).

#### Estimating the Motion Model 〈*P*〉

It is typical in fluorescence microscopy to perform a background subtraction (Lindeberg, 1998; Murphy & Davidson, 2012) when dealing with images where signal detail does not blend with the background. However, this approach is insufficient for data sets that include moving fluorescent particles that are occasionally obscured by locally high background levels or stationary fluorescent particles (see below). Thus, we designed an algorithm that relies on differences in the noise statistics between models of moving particles and stationary noise to better identify particles of interest. We engineered a solution that utilizes the properties of Poisson-like statistics in *K* to identify particles from regions where the noise statistics suggest evidence of a moving particle.

To resolve the moving particles from background noise, we first created two models from the primary data, one representing the moving particles (e.g., fluorescently labeled kinesin) and the other representing stationary signals. The processively moving kinesin motors present in the effective photon count model *K*, can be readily followed over background by eye when visualizing the time sequence as a movie at 10 frames per second (fps). However, it is difficult to computationally identify these motile particles using an automated algorithm that processes frames independently. We sought inspiration from the human visual system (Lindeberg, 1998), because it correlates pixels in adjacent frames of a movie to augment object recognition. We performed a spatiotemporal convolution of the pixels in *K* according to a ballistic diffusion model to supplement the signal of dim photon signatures from moving motors. We reasoned that a pixel, *i*, at frame, *j*, should be temporally correlated to a Gaussian blurred pixel, *i*, at frame, *j* ± *n*, where the Gaussian filter has a standard deviation (σ) of σ ∝ *n*. Each pixel *i* at frame *j* was averaged with its temporal neighbors with weights calculated from a 1D Gaussian kernel with a standard deviation (σ), of 1 frame.

In the interests of computational efficiency, we truncated the length of our temporal kernel, *T*(*n*), to a size of 5 temporal pixels. We generated two Gaussian filter movies, one with a standard deviation (σ) of 1 pixel for pixels in adjacent frames to pixel *i* which we denote as *G*_1_(*K*) and one with a standard deviation (σ) of 2 pixels for pixels 2 frames before and after pixel *i* which we denote as *G*_2_(*K*). For every pixel *i* at frame *j* the estimated motion model 〈*P*〉 mean as:

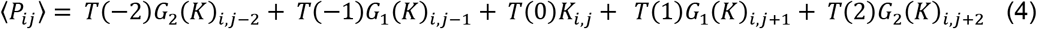

For this work, all of our convolution operations use reflective boundary conditions.

#### Estimating Stationary Model 〈*Q*〉

The program next generates a stationary model 〈*Q*〉 to represent the background fluorescence. The effectiveness of Cega is reliant upon how well the stationary model estimates the background of the motion model. Since we used a sequence of expanding Gaussian filters to estimate 〈*P*〉, a similar level of blurring is required for the estimate of 〈*Q*〉, or there will be structural artifacts wherever the background has sharp features. To ensure 〈*Q*〉 has the same resolution as 〈*P*〉, we generate an intermediate movie of convolved filters without temporal correlations.

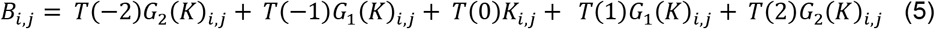

The temporal median filter is then applied to every pixel in *B*. We chose a sliding window of 31 frames, 15 frames before and after the pixel of interest to sample our median pixel as it appeared to provide the best compromise between dynamics suppression and background estimation accuracy. Ideally, the median filter needs to suppress dynamical fluctuations from moving motors while representing a gradually fluctuating background as accurately as possible. 〈*Q*〉 is defined as the movie generated after applying a non-temporal averaging of the Gaussian filters used on 〈*P*〉, followed by a temporal median filter with a sliding window of 31 frames.

#### KL Divergence - Segmenting Moving Particles from Noisy Data

Next, the differences between moving 〈*P*〉 and stationary 〈*Q*〉 models are used to resolve moving particles from stationary noise. To do so, we used the Kullback-Leibler (KL) divergence (Kullback & Leibler, 1951) to estimate the dissimilarity between the stationary and motion models. The general approach is to estimate two separate model hypotheses from the movie *K* and apply the KL divergence on a per-pixel basis that calculated the dissimilarity between groups.

The KL divergence for some random quantity *x* given hypothesis *P* and tested against null hypothesis *Q* is defined as:

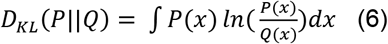

In our application, we posited two hypotheses with Poisson statistics, the motion 〈*P*〉 and stationary 〈*Q*〉 models, where the quantity *x* represents the effective photon counts that could be measured in an image pixel. We hypothesized that the movie *K* is a single instance of *x*, a Poisson-like realization of the per pixel distribution *P*. We wanted to see how poorly a per pixel distribution *Q* would predict *K* if *P* more accurately modeled the input data and *Q* only modeled stationary objects. In other words, either the pixels in 〈*P*〉 or 〈*Q*〉 can be used to represent the expectation value of the pixels in *K*, but since they are different models they will lead to different probability densities per pixel. Since *K* is Poisson-like, this means that the KL divergence of the *i*-th pixel in *Q* from the *i*-th pixel in *P* is the KL divergence between two Poisson distributions:

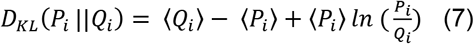

Given estimators for 〈*P*〉 and 〈*Q*〉, the KL Divergence movie (KLM) is generated.

#### Connectivity Filtering - Denoising the KL Divergence Movie

The KLM has larger pixel values where the two models are in disagreement, suggesting evidence of mobile motors since 〈*Q*〉 was designed to ignore motion. However, the estimators are not perfect, and this results in unusually noisy pixels throughout the movie where 〈*P*〉 and 〈*Q*〉 are in spurious disagreement with one another. This produces white pixels, known as salt noise, that are sprinkled in throughout the movie, without producing darker pixels, or pepper noise. Since the data is under sampling the Nyquist limit, the additional effect of temporal averaging in 〈*P*〉 all but guarantees that the signal left by the moving motors spreads over multiple pixels. The spurious noise of the KLM is removed with a connectivity filter. To do this, any *3×3* pixel subregion in the KLM must have at least 3 pixel values greater than 0.1 nats (units of natural logarithm), or the center pixel of that subregion is set to 0 in the connectivity movie.

#### LoG Filtering and Detecting Local Maxima

The denoised KLM is then passed through a scale space Laplacian of Gaussian (LoG) filter (Lindeberg, 1998) with two sigma values, 1 and 1.5 pixels, representing the parameter width of the filtering kernels, to generate the LoG movie. The smaller sigma value represents a typical motor signal and the larger sigma value helps to identify potential motors that are moving much faster and display a smeared information signature in the denoised KLM. We did not notice an improvement in object detection with additional scales in the LoG filter. Other SPT programs such as u-track (Jaqaman et al., 2008) and TrackMate (Tinevez et al., 2017) also use filters based on LoG segmentation, but apply them to raw data.

Local maxima from the LoG filtered movies are then aggregated into a list of potential signal coordinates. The coordinates for each local maximum (x, y, t) are used to retrieve the KL divergence score of the associated maximum pixel. Coordinates that point to a *3×3* region of KL divergence values that sum to less than a user-specified threshold are discarded.

### Characterization/Optimization of Performance

We used Cega to analyze fluorescently labeled kinesin motors moving along native cellular microtubule networks preserved after extracting membranes and soluble cytosolic components (Figure 1 A; see Methods). These data have substantial background fluorescence along with a high proportion of stationary fluorescent particles due to nonspecific binding that together represent experimental noise. Movies of a truncated GFP-kinesin-1 construct (GFP-K560) moving along microtubule arrays in axonal and dendritic compartments of extracted rat hippocampal neurons were acquired with an EMCCD camera where *R* < 1 effective photon, *G* = 70 gain and *O* = 2993 offset (see Methods for estimation). In axons, kinesin-1 motors move unidirectionally outward from the cell body, due to the unipolar arrangement of the microtubule cytoskeletal tracks. However, in dendrites, kinesin-1 motors move in both outward (anterograde) and inward (retrograde) directions due to the mixed polarity of the microtubule cytoskeleton in this part of the neuron. Our goal was to better understand the characteristics of the motile behaviors of kinesin in these environments. However, manual analysis and current SPT software could not reliably distinguish moving motors from nuisance particles in the background. Thus, we developed and optimized Cega to analyze these data sets.

The first step in implementing Cega to resolve moving kinesin motors from noisy background fluorescence was to perform pixel calibration (Figure 1B and Table 1). The calibrated model we calculated was similar to the initial uncalibrated data after brightness and contrast enhancement, and maintained the signal from both moving and stationary particles as well as the background noise (Table 1 and Figure 1C, Calibrated). We then used this calibrated model to generate both the stationary and motion models (Table 1 and Figure 1C, Motion and Stationary). Each frame of the resulting motion and stationary movie resembled a softened version of the calibrated movie. The number of frames used to generate the stationary movie was larger than that of the motion movie, and resulted in the same image over the three time points sampled in Figure 1C. Most of the signal from moving motors was eliminated in the stationary model, leaving background noise. In contrast, the motion movie contained signal from moving motors and background. Then, we used the KL-divergence to estimate the dissimilarity between the stationary and motion models, forming the KL-divergence movie (Table 1 and Figure 1C, KL-divergence). The KL-divergence model eliminated the background noise while enhancing the signal of moving motors. The reader can see that this step clearly isolates the particles of interest over the background. However, this step introduced noise in areas where the stationary and motion models are in disagreement; this noise was eliminated with a connectivity filter (Table 1 and Figure 1C, Connectivity). The signal that remained represented moving particles (colored circles in connectivity row; Figure 1C). The connectivity movie was then passed through a scale space Laplacian of Gaussian (LoG) filter (Lindeberg, 1998) to enhance the edges of the signal left from the connectivity filter (Table 1 and Figure 1C, LoG). This step generated signal surrounding the moving particles, representing their boundaries (colored circles in LoG row; Figure 1C). Local maxima from the LoG filtered movies were then aggregated into a list of potential signal coordinates (Table 1, LoG Filter). The corresponding KL-divergence score for each of the local maxima was used to threshold potential particle candidates. Using tracking software, these coordinates were connected into trajectories.

**Table 1.**
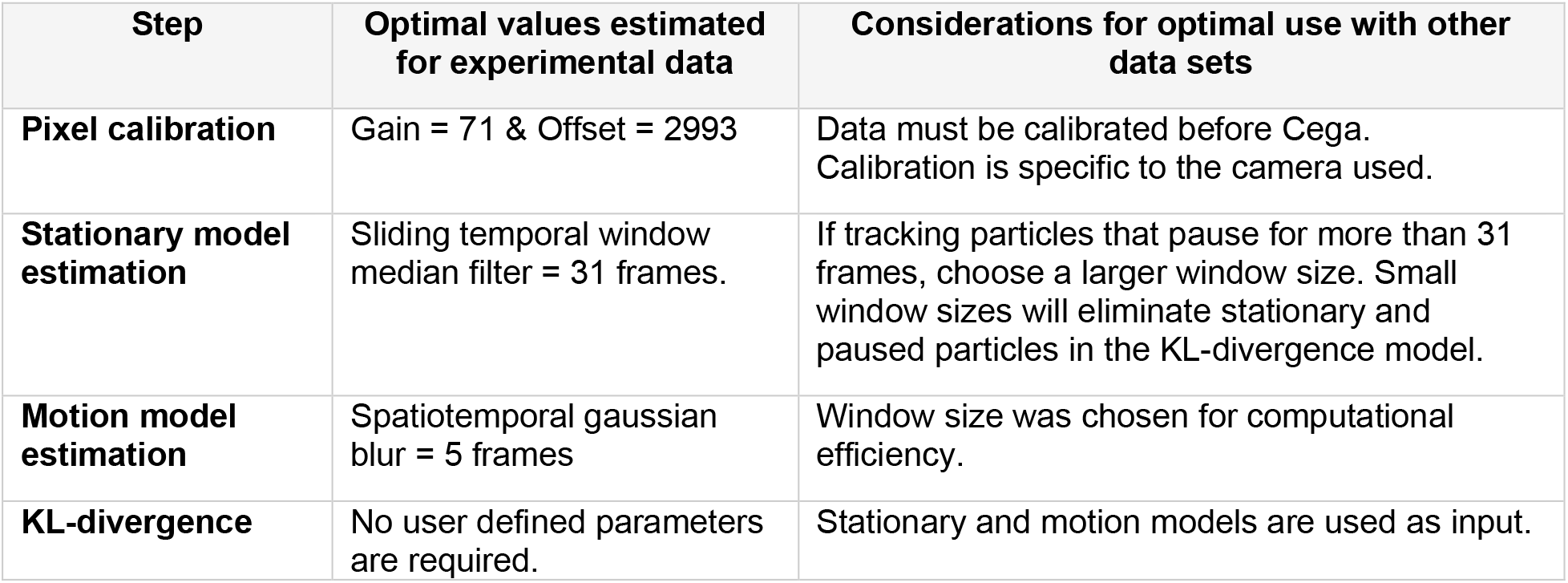
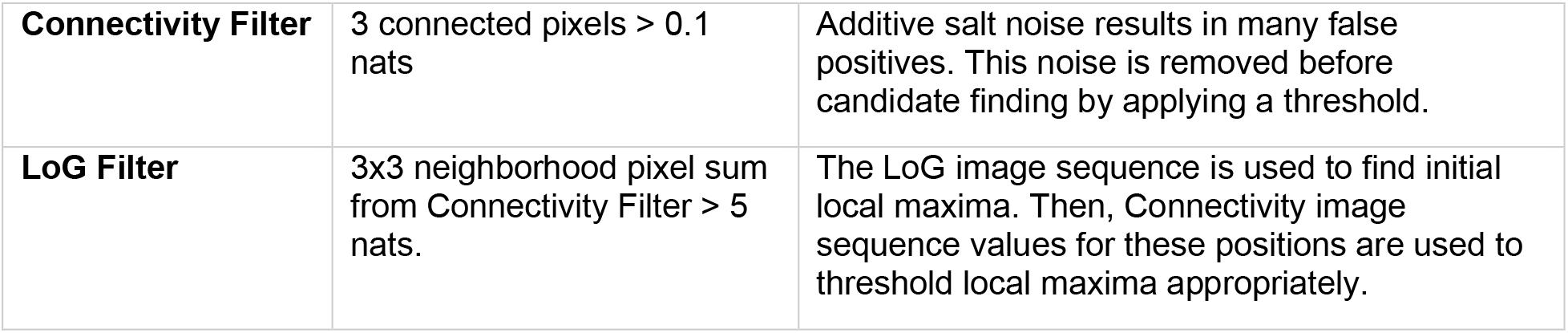
Cega thresholding values used for experimental data.

| **Step** | **Optimal values estimated for experimental data** | **Considerations for optimal use with other data sets** |
| --- | --- | --- |
| **Pixel calibration** | Gain = 71 & Offset = 2993 | Data must be calibrated before Cega. Calibration is specific to the camera used. |
| **Stationary model estimation** | Sliding temporal window median filter = 31 frames. | If tracking particles that pause for more than 31 frames, choose a larger window size. Small window sizes will eliminate stationary and paused particles in the KL-divergence model. |
| **Motion model estimation** | Spatiotemporal gaussian blur = 5 frames | Window size was chosen for computational efficiency. |
| **KL-divergence** | No user defined parameters are required. | Stationary and motion models are used as input. |
| **Connectivity Filter** | 3 connected pixels > 0.1 nats | Additive salt noise results in many false positives. This noise is removed before candidate finding by applying a threshold. |
| **LoG Filter** | 3x3 neighborhood pixel sum from Connectivity Filter > 5 nats. | The LoG image sequence is used to find initial local maxima. Then, Connectivity image sequence values for these positions are used to threshold local maxima appropriately. |

### Comparison to Existing Methods

#### Candidate Finding

To test Cega’s ability to correctly detect simulated particles over typical background noise, we generated a time series of simulated particles by adding simulated photons on temporally filtered experimental data (see simulation methods). Fluorescent particles that represent GFP-K560 were simulated with mean photon emissions ranging from 50 - 600 photons per full frame of acquisition. This range includes the peak photon range of motor signal within our experimental data, which was measured to be between 100-200 photons following integration time. Moving particles were detected using Cega and background subtraction of median or minimum temporal filters with a 31frame window, or standard particle detection with no background subtraction (Lindeberg, 1998). All methods were manually tuned with thresholds to maximize performance at a specific probe intensity, but the background was always the same. The spot finding algorithms had a threshold based upon the maximum pixel value in a 3×3 pixel neighborhood of the estimated spot center. To quantitatively compare Cega’s performance against other particle detection methods processes, we calculated the Jaccard index and recall rate (see Methods). The Jaccard index is a statistic used to understand the similarities between sample sets (Milligan, 1981), and in this case is a measure related to the similarity between spot positions detected by each of the detection processes compared to the actual simulated spot positions. The recall rate is the sensitivity between spot positions detected by each of the detection processes compared to the actual simulated spot positions.

The performance of Cega was ranked amongst typical spot finding algorithms that use the raw movie or a background subtracted movie as the signal prior to scale space LoG filtering (Figure 2). No algorithm tested was capable of providing a tracking solution at 50 photons, but Cega showed noticeable improvements at 100 photons and the median background subtracted spot finder matched performance after 200 photons, which is greater than the range of our experimental data. The minimum background subtracted spot finder met up with the prior two algorithms at 350 mean photons. The standard spot finder did not catch up in performance but started to improve significantly after 350 photons.

**Figure 2.**
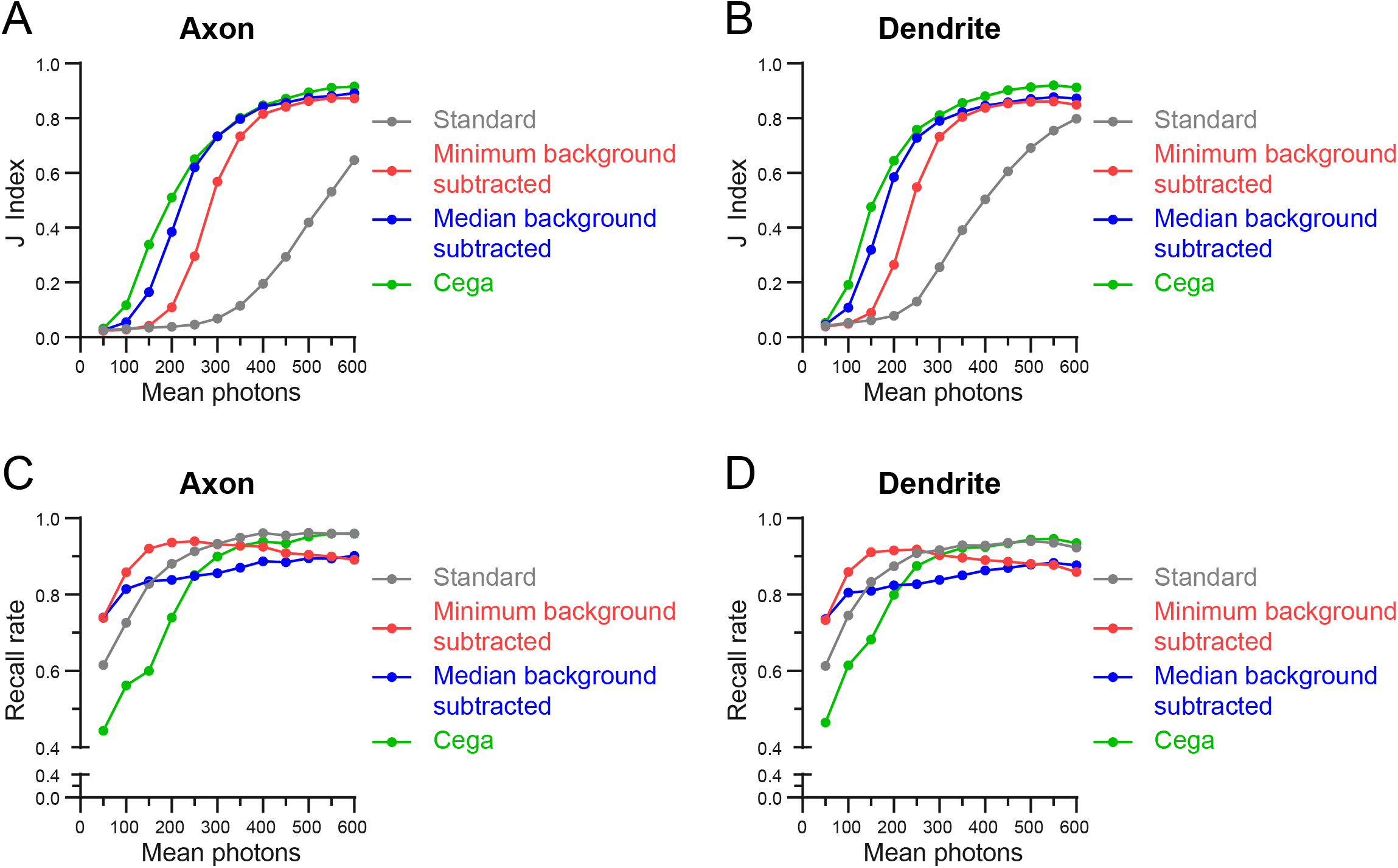
Analysis of Cega spot detection. A and B) Jaccard index values calculated for simulated spots detected with Cega, median or minimum background subtraction methods, or standard methods, as mean photon count is increased. C and D) Recall rate for detection methods as mean photon count increases.

The trends from the Jaccard index test show that any sort of background subtraction is a vast improvement over spot finding with the raw image (Figure 2A and B). The trends also show that Cega is able to discern between false signal and a moving motor with less information than the background subtracted methods. The recall rate was high for all methods at high probe intensities but was at 50% for Cega at 50 photons (Figure 2C and D), since Cega could not reliably extract particle features at this photon count. The other methods which had twice the recall rate at 50 photons had similar or lower Jaccard indices, which meant they had twice as many false positives. Finer parameter tuning could raise or lower Jaccard indices by a few percentage points, but the trends are clear, Cega performs more reliably at lower probe intensities and background subtraction methods converge in performance shortly thereafter.

#### Tracking

We also tested the ability of Cega to effectively discard stationary particles, which becomes apparent when performing a full tracking experiment. After candidate finding, simulated motor spot coordinates were connected into trajectories. The trajectories formed after application of Cega or median background subtraction were then displayed on kymographs and compared against the known simulated spot trajectories (Figure 3). Kymograph plots of simulated tracks show that Cega provided for more accurate tracking of simulated particles, compared to median background subtraction which had lower Jaccard indices. The zoom-in overlap shows that the median background subtraction methods resulted in many false tracks, most likely due to capturing background noise. Comparison of the Jaccard index and recall rates showed that Cega was more accurate in generating known tracks. This was true for simulated movies from both axons and dendrites with differing background signal. The difference in Jaccard index between tracks resulting from Cega and median background subtraction was greatest for the simulated axonal movie which contained uneven illumination and more background noise than the dendritic movie. In addition, Cega allowed for improved tracking in the simulated dendrite movie, where particle density was higher.

**Figure 3.**
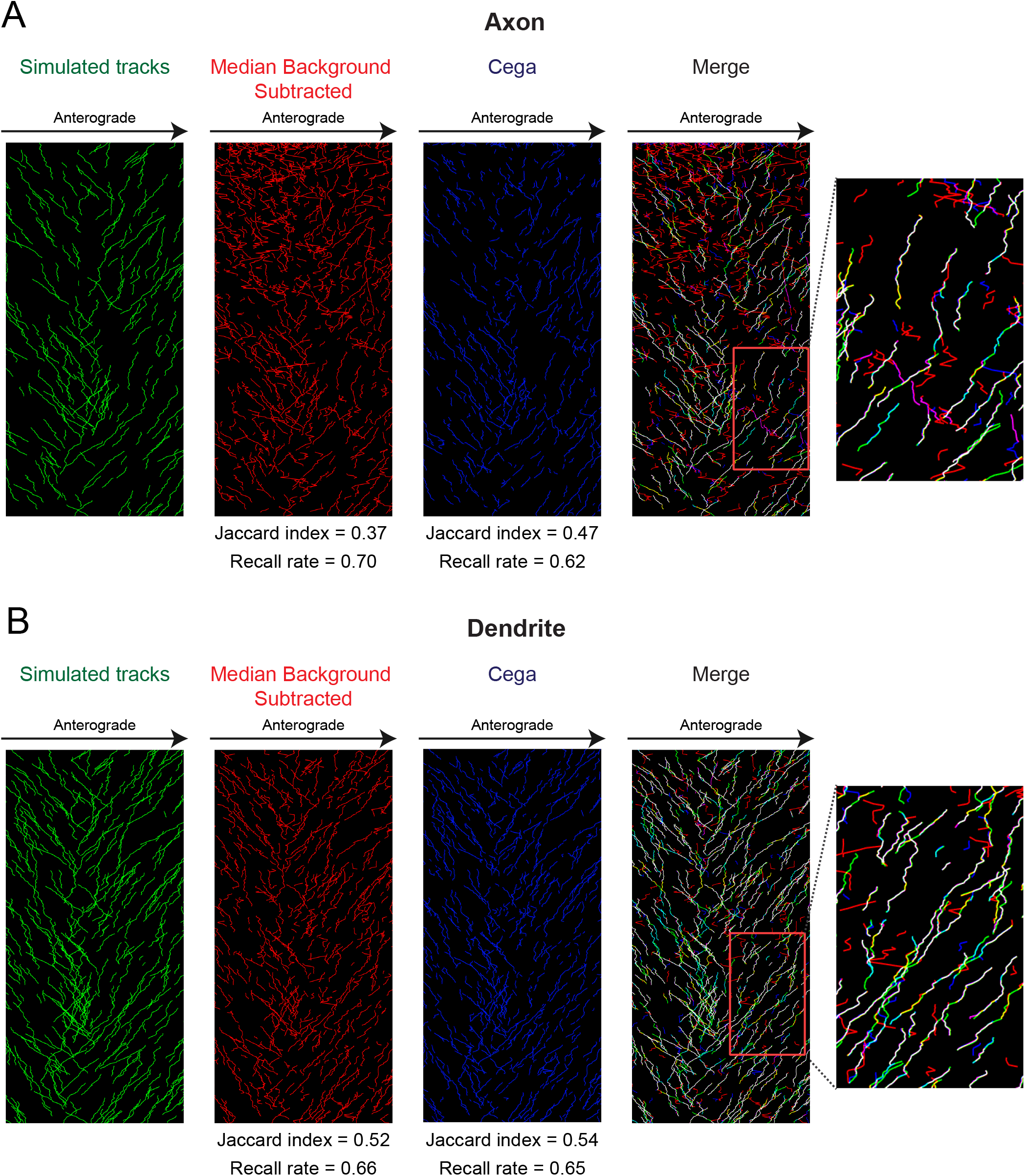
Analysis of tracking after Cega or median background subtraction methods. A and B) Kymographs of tracks determined from simulated particles within axonal and dendritic compartments. Although the same number of particles were simulated in the axonal and dendritic movies, the dendritic movie was smaller in size, resulting in a higher density of particles. In the merge kymograph and zoom-in area, cyan indicates locations where only simulated tracks and Cega detected tracks overlap, whereas yellow indicates where only simulated tracks and median background subtracted detected tracks overlap. Magenta tracks are where only Cega and median background subtracted tracks overlap. Jaccard indices and recall rates for tracks determined after Cega filtering and median background subtraction are listed below each corresponding kymograph.

After testing Cega with simulated data, we used Cega to analyze our experimental data. Motor tracks detected during Cega analysis of experimental movies were mostly linear, with velocities similar to those expected from kinesin-1 motors in an *in vitro* system, while median background subtraction produced many tracks containing spurious noise, resulting in non-uniform tracks. We also found that Cega was able to track motors that switched direction during movement (Figure 4B zoom-in). As these directional switches were not detected in this data set with programs such as u-track (Jaqaman et al., 2008) and TrackMate (Tinevez et al., 2017), the Cega algorithm provides a more reliable analysis of the data from these complex microtubule arrays.

**Figure 4.**
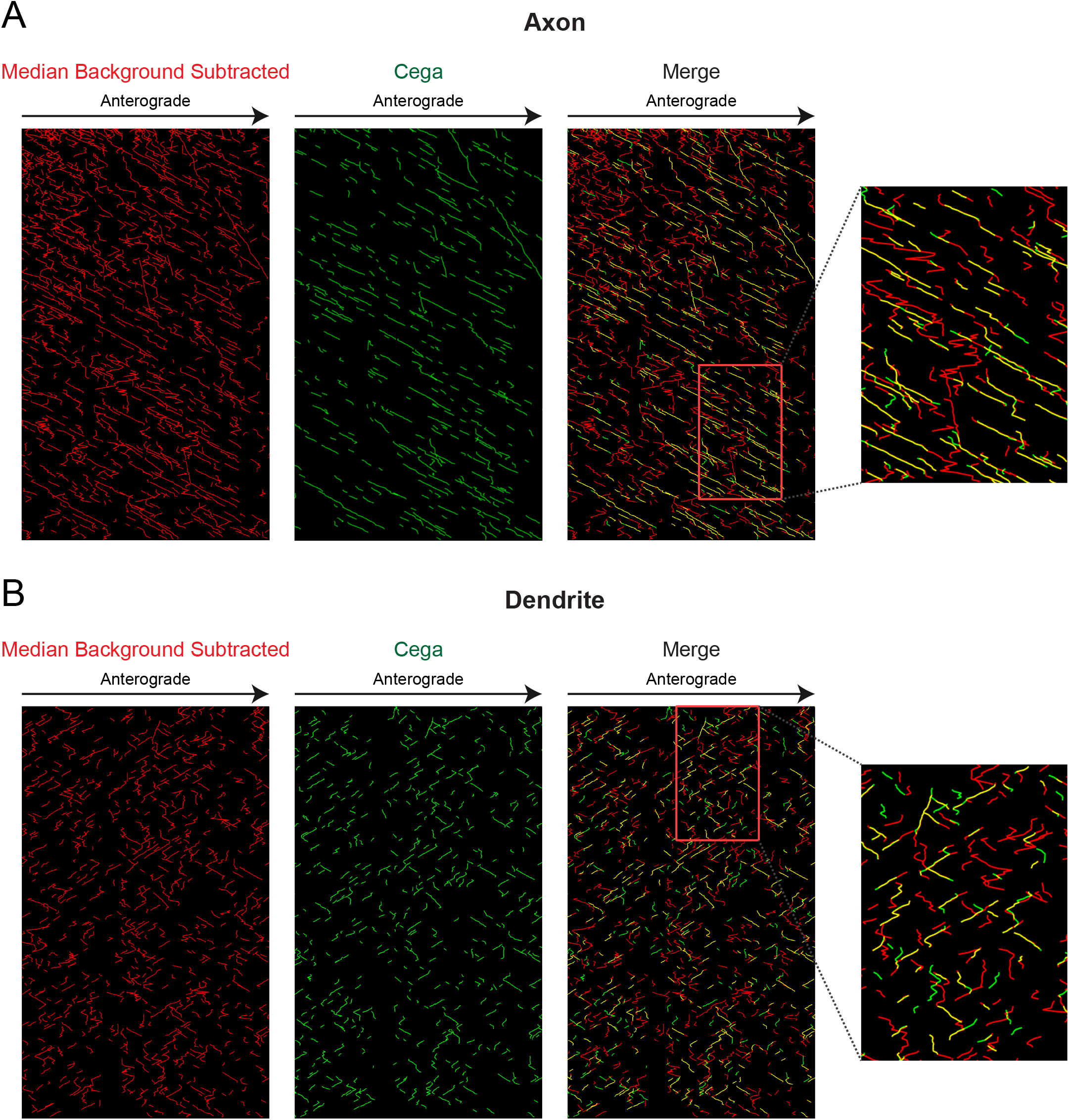
Analysis of Cega performance on original dataset. A and B) Kymographs of tracks determined from kinesin motors moving on microtubule arrays within axonal and dendritic compartments. In the merge kymograph, yellow areas indicate where Cega and median background subtracted detected tracks overlap.

## DISCUSSION

Single particle tracking software allows automated quantitative analysis of both *in vitro* and *in vivo* experimental data. However, as experimental systems become more complex, background noise often increases, posing a significant challenge to current software. As an example, we analyzed the motility of kinesin motors moving along extracted neuronal cytoskeletal arrays, as these data are much noisier than data from more reductionist single molecule *in vitro* assays. The quantitative metrics and qualitative comparisons shown here make it clear that more sophisticated particle identification methods are required to reliably extract SPT information from noisier data sets. Conventional spot finding algorithms use pixel intensity metrics for thresholding of real signals from artifacts. In our datasets from motility along extracted neurons, intensity-based metrics failed to reliably threshold real motor signals from noise. As a result, more than half of the trajectories recovered from SPT software were false. Cega, on the other hand, uses statistical calculations which factor in background estimations so that artifacts from local regions of high background fluorescence are mitigated. Use of Cega on experimental data resulted in a dramatic reduction and temporal shortening of false trajectories.

We found that Cega was dependent on the quality of the background estimation technique used to determine 〈*Q*〉. Before settling on a temporal median background estimator, we first tried using the temporal minimum filter, but we found that the median filter was a better discriminator of signal over noise at lower photon counts. It is possible that there is an ideal quantile value for background estimation, but it was not an avenue we explored because it seemed that finding the optimal quantile would be dependent on specific experimental conditions. Another avenue for optimization would be to incorporate a weighted spatiotemporal median filter instead of the standard temporal median filter (Brownrigg, 1984). The current implementation of Cega out-performs current background subtraction methods when analyzing neuronal data sets. Still, we believe that further improvement of Cega is feasible and future studies should focus on improving the accuracy of background estimation.

The Jaccard index and recall rate from a range of detection techniques were compared to demonstrate that Cega had better performance than a typical spot finder with background subtracted data (Hoogendoorn et al., 2014). In either instance, the recall rate of both methods could approach 100%, but the number of false positives would skyrocket; this information on spot reliability is conveyed by the Jaccard index. Unfortunately, the Jaccard index does not fully represent tracking capability, as a false negative compromises tracking software much more than a false positive (Smal & Meijering, 2015), but this is only true when false positives are not spatiotemporally correlated to one another. We treat the Jaccard index as a baseline for the potential of a successful tracking analysis. A Jaccard index over 0.9 is very likely to return accurate, full length trajectories. A Jaccard index of 0.5 is more likely to return partial trajectories if the recall rate is low or is more likely to return a high percentage of false trajectories if the recall rate is high and the artifacts are correlated. Therefore, although the Jaccard index and recall rate alone are not sufficient to guarantee tracking performance, together they provide good indicators as to assess the relative quality of tracking algorithms.

Our simulations were designed to be optimal for our data sets. The simulations were generated by adding photons from simulated motors on a median background estimate of axonal and dendritic movies. The median background was taken from a much shorter temporal window (7 frames) than what is used from our spot finding analyses (31 frames). The reason for this was that a shorter temporal median filter preserved the background structure better. Legitimate motors from experimental data were not completely filtered out as a direct consequence of using a shorter temporal window in the median filtering of our simulated data. As a result, there were spatiotemporally correlated pixels that highlight moving particles in our background data. The estimation artifacts guaranteed that none of the measured spot finding algorithms could score a perfect Jaccard index of 1. The moving background artifacts resulted in a few spurious particle coordinates in Figure 2 which actually corresponded to true trajectories in Figure 3. Nevertheless, all spot finding algorithms we tested tracked these artifacts.

SPT remains an open problem and the field is constantly evolving. For this work, we used a consistent tracking software so we could focus on assessing the quality of the initial spot detection algorithm chosen. The in-house tracking software used here, based on the linear assignment problem (LAP) used in u-track (Jaqaman et al., 2008), in combination with Cega, allowed for tracking of moving motors within a dense background. Importantly, Cega spot detection allowed for the tracking of motors that change directions, something that programs such as u-track (Jaqaman et al., 2008) and TrackMate (Tinevez et al., 2017) were unable to accomplish with this experimental dataset. However, the tracking software we used is still not optimal for this problem; aside from the spot candidates, the additional information generated from Cega is not incorporated into the downstream tracking algorithms. Incorporating the heterogeneous background information or the KL divergence score into downstream tracking methods would increase the robustness of the tracker in noisier data. Most notably is the localization algorithm used in this study. Our chosen tracker has a localization algorithm incorporating MAPPEL software (Olah, 2017/2019), which does not incorporate the estimated background information from Cega. As a result, the returned fit statistics are sub-optimal indicators for the tracking phase. It is well known that incorporating prior background information greatly enhances the accuracy recall rate of single molecule microscopy (Hoogendoorn et al., 2014), so measuring the trajectory improvements from an enhanced localization estimator would be a good next step in improving tracking resources for motors community.

Although there are many components in SPT that are yet to be refined for tracking in more complex data sets such as the neuronal data analyzed here, Cega presents a marked improvement in a critical component of the greater SPT workflow. By using statistical hypothesis techniques to segment background, we were able to track moving particles more reliably while discarding nuisance objects. Our refined trajectories dramatically minimize false trajectories because Cega is a better discriminator of signal than intensity-based spot finding algorithms. By estimating a baseline of accuracy in the beginning phases of a tracking algorithm, we can dramatically improve downstream performance for all SPT algorithms that are structured to identify particles before the connecting trajectories.

## METHODS

### Protein Expression and Purification

K560-GFP protein was purified as described in (McIntosh et al., 2018) with the following modifications. The K560-GFP DNA was transformed into BL21(DE3)pLysE bacteria (Sigma Aldrich, CMC0015-20X40UL) and cultures containing the plasmid were grown at 37°C until an OD600 of 0.4 was reached. Protein expression was then induced with 0.15 mM IPTG for 18 hrs at 18°C. Cells were pelleted and flash frozen in liquid nitrogen and stored at −80°C. On the day of purification, cells were resuspended in lysis buffer (50 mM NaPO4, 250 mM NaCl, 20 mM Imidazole, 1 mM MgCl2, 0.5mM ATP, 1 mM $\beta$-ME, 0.01 mg/mL aprotinin and leupeptin, pH 6.0), and lysed by passage through a microfluidizer (Microfluidics). Lysate was clarified by centrifugation at 42,000 rpm for 30 minutes, and subsequently run over a Co2+ agarose bead (GoldBio, H-310-25) column at 1 mL/minute. Bound protein was washed with wash buffer (50 mM NaPO4, 300 mM NaCl, 10 mM Imidazole, 1 mM MgCl2, 0.1 mM ATP, 0.01 mg/mL aprotinin and leupeptin, pH 7.4), and eluted with elution buffer (50 mM NaPO4, 300 mM NaCl, 150 mM Imidazole, 1 mM MgCl2, 0.1 mM ATP, pH 7.4). Elution fractions were pooled and concentrated. Buffer was exchanged to BRB80 (80 mM NaPIPES, 1 mM MgCl2, 1 mM EGTA, pH 6.8) by loading protein over Nap10 (GE Healthcare, 17-0854-01) and PD10 (GE Healthcare, 17-0851-01) exchange columns. MT affinity/dead-head spin was performed as described in (McIntosh et al., 2018). Protein concentration was determined using Pierce BCA Protein Assay Kit (ThermoFischer Scientific, 23225).

### Neuronal Cell Culture

35-mm glass-bottom dishes (MatTek, P35G-1.5-14-C) were coated with 0.5 mg/ml poly-L-lysine (Sigma-Aldrich, P1274) overnight, and rinsed with dH20 and MEM (Gibco®, 1109-072) prior to plating neurons. E18 Sprague–Dawley rat hippocampal neurons were received from the Neuron Culture Service Center at the University of Pennsylvania and plated in attachment media (MEM(Gibco®, 1109-072) supplemented with 10\% horse serum (Gibco®, 26050-070), 33 mM glucose (Corning, 25-037-CIR), and 1 mM sodium pyruvate (Gibco®, 11360-070)) at a density of 250,000 cells per dish and cultured 37°C with 5% CO2. After 4-6 hours of attachment, media was replaced with maintenance media (Neurobasal (Gibco®, 21103-049) supplemented with 33 mM glucose (Corning, 25-037-CIR), 2 mM GlutaMAX (Gibco\R, 35050-061), 100 units/mL penicillin and 100 µg/ml streptomycin (Gibco®, 15140-122), and 2% B27 (Gibco\R, 17504-044)). 24 hours after initial plating, AraC (Sigma Aldrich, C6645) was added at 10 μM to prevent glial cell division.

### Motor-PAINT Assay

The neuronal MT cytoskeleton was extracted, stabilized and fixed according to previous work (Tas et al., 2017) with slight modifications. At 9-10 DIV, membranes were extracted from neurons by incubation with extraction buffer (1M sucrose, 0.15% Triton-X in BRB80 pH 6.8 at 37°C) for 1 minute. An equal amount of fixation buffer (1% paraformaldehyde (PFA) in BRB80 pH 6.8 at 37°C) was added for 1 minute with gentle swirling. Dishes were rinsed 3 times with wash buffer (2uM Paclitaxel in BRB80 pH 6.8 at 37°C), and once more right before imaging. For imaging, extracted MT arrays were placed in imaging buffer (1 mM ATP, 2 uM Paclitaxel, 0.133 mg/mL casein (Sigma Aldrich, C5890-500G), 0.133 mg/mL bovine serum albumin (BSA) (Fischer Scientific, 50-253-90), 4 mM DTT, 6 mg/mL glucose, 49 U/mL glucose oxidase (Sigma, G2133-250KU), 115 U/mg catalase (Sigma, C100-500MG), 0.21 mg/mL creatine phosphokinase (Sigma, C3755-35KU), 4.76 mM phosphocreatine (Sigma, P7936-1G) in BRB80 pH 6.8) containing 5-10 nM K560-GFP motor dimer.

### Microscopy

Motor-PAINT assays were performed at 37°C using a PerkinElmer Nikon Eclipse Ti TIRF system, using a Nikon Apo TIRF 100x 1.49 NA oil-immersion objective and a Hamamatsu ImagEM C9100-13 EMCCD camera operated by Volocity software. Movies were obtained by continuously acquiring K560 motor images at 5 frames/second for 2 minutes.

#### Offset and Gain Calibration

Gain calibration of the Hamamatsu ImagEM C9100-13 EMCCD camera was performed by fusing techniques from standard and automated gain calibration practices. Our camera had a few damaged pixels in the upper right quadrant of the sensor Region of Interest (ROI). The damaged pixels caused errors in automated gain calibration (Heintzmann et al., 2018) but we performed some modifications to existing software to return reasonable calibration parameters. We were able to reliably track molecules with scalar gain, offset, and read noise variance parameters by cropping the sensor ROI so that only undamaged pixels were used in the following gain regression algorithms.

First, a dark movie was acquired with 200 frames with the camera shutter closed. The mean pixel value of the dark movie used the estimate for *O* and the corresponding variance was the estimate for *var*(*R*) in units of *ADU*^2^.

We performed the single shot gain estimator on a few cropped images given knowledge of *O* and *var*(*R*) and we extracted an averaged *G* as our gain parameter using the single shot gain estimation algorithm described in (Heintzmann et al., 2018). With *G* we were able to determine that *var*(*R*)*G*^−2^ ≪ 1 on this camera which allowed us to ignore the effects of additive read noise for the remainder of our analysis. The technique failed when processing with the complete camera ROI because a strip of about 6 adjacent camera pixels that appeared to have been damaged by a photon over-saturation event resulted in an over estimation of the energy contribution from the high frequency Fourier spectrum. The damaged pixels were noticeably darker than the more functional neighboring pixels, but they were spatially correlated, limited in number and did not noticeably affect tracking results or biological experiments.

We estimated our Poisson movie *K* ≈ (*S* − *O*)*G*^−1^ and discarded the read noise term. We use the movie *K* for all of our subsequent analysis for Cega and downstream tracking.

#### Motor Protein Simulations

Simulated data was generated by fusing a background estimate of experimental data with simulated photon signatures from procedurally generated trajectories. The resulting process provided a realistic movie with simulated motor photons and a ground truth of averaged positions.

The experimental movie was stripped of its true motor proteins via a median filter with a temporal width of 7 frames. The shortened temporal median filter provided a more accurate depiction of the experimental background data but left some noticeable residual emissions from moving motors wherever the motors had aggregated. This resulted in data where a certain fraction of false positives was guaranteed with experimentally relevant photon intensities for the simulated motors. Poisson noise was added to the median filtered movie.

The motor photon trajectory positions were calculated at time points before and after every simulated camera acquisition. The rough positions were modeled from a Kalman filter process, where proteins are subjected to diffusion with drift, but a white process noise is applied to the drift velocity so that proteins could gradually change direction and speed. The initial mean velocities were drawn from a log normal distribution to match the distribution of velocities observed in experimental data. Motor protein photons were painted by interpolating discrete trajectory positions with a Brownian bridge (Lindén et al., 2016) given exponentially sampled time points with a rate set by the mean photon parameter.

The final simulation movie consists of the simulated motor photons, summed with the background movie and white noise. The white noise had a sigma of 0.1 photons was added to all pixels to simulate extra read noise.

#### Jaccard Index Score and Recall Rate

The Jaccard index (Milligan, 1981) is an algorithm performance metric defined as:

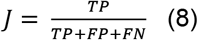

Where *TP* is the number of true positives, *FP* is the number of false positives, and *FN* is the number of false negatives when comparing a list of coordinates to a known ground truth. Similarly, the recall rate is defined as:

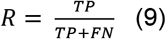

and is useful for determining how well a true trajectory can be reconstructed if there was a downstream method for removing false positives.

To score the performance of Cega and the other spot finders, the linear assignment problem (Jonker & Volgenant, 1987) was used to assign the nearest true coordinates to the algorithm coordinates. The squared distance between ground truth and algorithm coordinates plus 1 was used as the cost for assignment. The birth and death costs were set to 4. Any assignment costs greater than 4 were discarded to prevent assigning coordinates pairs that were separated by at least 2 pixels. Coordinate pairs that were assigned to one another were counted as true positives, true coordinates did not pair were scored as false negatives, and algorithm coordinates that did not pair were scored as false positives. For trajectories formed after spot detection with Cega and the other spot finders, the Jaccard index and recall rates were calculated by taking each kymograph and measuring the overlap of track segments to the ground truth on a pixel basis.

#### Tracking Software

The tracking software implemented for this manuscript was adapted from the MATLAB software developed for (Schwartz et al., 2017). A few minor modifications were applied to make the workflow amenable to motor proteins, primarily the blob finding routine for initial candidate selection was replaced with Cega. The software in its current implementation does not yet incorporate heterogeneous background information in the localization routines. A median background subtraction on the data was considered for the localization routine, but was omitted from this exposition because the maximum a Posteriori estimator cannot process negative pixel values. The same tracking software and parameters were used for all comparisons between Cega and the median subtracted spot finder.

## ACKNOWLEDGMENTS

This work was supported by the Center for Engineering MechanoBiology NSF Science and Technology Center, CMMI:15-48571, and National Institutes of Health grants RM1 GM136511 (to E.M.O, E.L.F.H and M.L.) and R35 GM126950 (to E.L.F.H).

## VIDEO LEGENDS

**Video 1. Cega processing steps for axonal movie.** Movies are displayed from top to bottom as follows: original movie, calibrated movie, stationary movie, motion movie, KL-divergence movie, connectivity filter movie, and LoG movie. Brightness and contrast were adjusted differently for each movie to show best comparison. Movie set to play 10 frames per second, and scale bar set at 10 μm.

**Video 2. Zoom-in of Video 1.** Green arrows indicate positions of moving particles while red arrows indicate positions of stationary particles. Movie set to play 10 frames per second, and scale bar set at 2 μm.

**Video 3. Cega processing steps for dendritic movie.** Movies are displayed from top to bottom as follows: original movie, calibrated movie, stationary movie, motion movie, KL-divergence movie, connectivity filter movie, and LoG movie. Brightness and contrast were adjusted differently for each movie to show best comparison. Movie set to play 10 frames per second, and scale bar set at 10 μm.

**Video 4. Zoom-in of Video 3.** Green arrows indicate positions of moving particles while red arrows indicate positions of stationary particles. Movie set to play 10 frames per second, and scale bar set at 2 μm.

**Video 5. Detected particles using Cega for each processing step of axonal movie.** Movies are displayed from top to bottom as follows: original movie, calibrated movie, stationary movie, motion movie, KL-divergence movie, connectivity filter movie, LoG movie, track movie with LoG movie overlay, cumulative track movie. All movies except the last consist of the original movie with red or green boxes added to indicate locations of detected particles. Red boxes indicate locations of detected particles for each filtering step, while overlaid green boxes indicate particle positions detected after LoG filtering with Cega. The track movie overlaid on the LoG movie shows tracks detected within the last 5 frames, while the cumulative track movie indicates all tracks detected up until the current frame. Movie set to play 10 frames per second, and scale bar set at 10 μm.

**Video 6. Zoom-in of Video 5.** Movie set to play 10 frames per second, and scale bar set at 2 μm.

**Video 7. Detected particles using Cega for each processing step of dendritic movie.** Movies are displayed from top to bottom as follows: original movie, calibrated movie, stationary movie, motion movie, KL-divergence movie, connectivity filter movie, LoG movie, track movie with LoG movie overlay, cumulative track movie. All movies except the last consist of the original movie with red or green boxes added to indicate locations of detected particles. Red boxes indicate locations of detected particles for each filtering step, while overlaid green boxes indicate particle positions detected after LoG filtering with Cega. The track movie overlaid on the LoG movie shows tracks detected within the last 5 frames, while the cumulative track movie indicates all tracks detected up until the current frame. Movie set to play 10 frames per second, and scale bar set at 10 μm.

**Video 8. Zoom-in of Video 7.** Movie set to play 10 frames per second, and scale bar set at 2 μm.

